# Deep learning prediction boosts phosphoproteomics-based discoveries through improved phosphopeptide identification

**DOI:** 10.1101/2023.01.11.523329

**Authors:** Xinpei Yi, Bo Wen, Shuyi Ji, Alex Saltzman, Eric J. Jaehnig, Jonathan T. Lei, Qiang Gao, Bing Zhang

**Affiliations:** Lester and Sue Smith Breast Center, Baylor College of Medicine, Houston, TX 77030, USA; Department of Molecular and Human Genetics, Baylor College of Medicine, Houston, TX 77030, USA; Department of Liver Surgery and Transplantation, Liver Cancer Institute, Zhongshan Hospital and Key Laboratory of Carcinogenesis and Cancer Invasion of the Ministry of China, Fudan University, 180 Fenglin Road, Shanghai 200032, China; Mass Spectrometry Proteomics Core, Advanced Technology Cores, Baylor College of Medicine, Houston, TX 77030, USA

## Abstract

Shotgun phosphoproteomics enables high-throughput analysis of phosphopeptides in biological samples, but low phosphopeptide identification rate in data analysis limits the potential of this technology. Here we present DeepRescore2, a computational workflow that leverages deep learning-based retention time and fragment ion intensity predictions to improve phosphopeptide identification and phosphosite localization. Using a state-of-the-art computational workflow as a benchmark, DeepRescore2 increases the number of correctly identified peptide-spectrum matches by 17% in a synthetic dataset and identifies 19%-46% more phosphopeptides in biological datasets. In a liver cancer dataset, 30% of the significantly altered phosphosites between tumor and normal tissues and 60% of the prognosis-associated phosphosites identified from DeepRescore2-processed data could not be identified based on the state-of-the-art workflow. Notably, DeepRescore2-processed data uniquely identifies EGFR hyperactivation as a new target in poor-prognosis liver cancer, which is validated experimentally. Integration of deep learning prediction in DeepRescore2 improves phosphopeptide identification and facilitates biological discoveries.

## Introduction

Post-translational modifications (PTMs) are ubiquitous in human cells and are broadly involved in the regulation of protein activity, localization, stability, and interaction. One of the most important and extensively studied PTMs is phosphorylation, which plays a critical role in regulating a wide range of biological processes and signaling pathways^1^. Shotgun phosphoproteomics, in which proteins are digested by proteases such as trypsin and analyzed via liquid chromatography tandem mass spectrometry (LS-MS/MS), provides an unbiased, high-throughput method to systematically study phosphorylation in biological samples^2^.

The first and absolutely essential step in phosphoproteomics data analysis is the identification of phosphopeptides based on MS/MS spectra, which is commonly achieved through database searching^3^. Multiple tools, such as MaxQuant^4^, MS-GF+^5^, X!Tandem^6^, Comet^6,7^, Mascot^8^, pFind^9^, SEQUEST^10^, and MSFragger^11^, are available for this analysis. These tools report identified phosphopeptide sequences along with assigned phosphorylation sites. However, the confidence of site localization is often not specifically determined. To determine the confidence of each possible phosphorylation site candidate in an identified peptide sequence, several computational algorithms have been developed. Mascot Delta score^12^ and SLIP^13^ compute the difference of peptide-spectrum match (PSM) scores associated with different competing phosphosites to determine site localization. PhosSA^14^ uses a dynamic programming algorithm for phosphorylation site assignment. Many other algorithms employ probability-based scores to determine phosphorylation site localization, such as AScore^15^ and phosphoRS^16^ implemented in PeptideShaker^17^, PTMScore^18^ implemented in MaxQuant, and PTMprophet^19^ implemented in philosopher^20^.

Despite the availability of numerous computational tools, phosphopeptide identification remains a challenging task, as clearly indicated by the high level of discrepancy among identification results generated by different computational tools on the same data^21^. Because each peptide sequence may include multiple candidate phosphorylation sites, the search space in phosphoproteomics is much larger compared with an unmodified search, and this can lead to reduced sensitivity and increased false positive rate in phosphopeptide identification and phosphosite localization. Phosphosite localization is further hampered by the lack of site-determining ions in many experimental spectra. As a result, the spectrum resolution rate in a phosphoproteomics experiment is much lower than that in a global proteomics experiment (**Figure S1**), limiting the potential of phosphoproteomics-based biological discoveries. Moreover, a low spectrum resolution rate in individual experiments further leads to many missing values in the resulting dataset, which further reduces the statistical power for biological discoveries.

Recent advancements in deep learning have provided new opportunities for proteomics^22^. Deep learning derived features, such as the retention time (RT) difference between experimentally observed and computationally predicted RTs and the similarity between experimentally observed and computationally predicted MS/MS spectra have been shown to effectively discriminate correct and incorrect PSMs in phosphoproteomics, and these features have been used as evaluation metrics to benchmark computational tools for phosphopeptide identification and phosphosite localization^21^. Incorporating these features into PSM rescoring has been shown to improve peptide identification in global proteomic profiling and immunopeptidomic profiling^23,24^,25. Recently, it has been shown that deep learning-based fragment ion intensity prediction can facilitate phosphorylation site localization^26^. However, the potential utility of retention time prediction in phosphorylation site localization has not been investigated. Furthermore, there is no method combining deep learning-facilitated PSM rescoring and site localization to improve the sensitivity and accuracy of phosphopeptide identification.

Built upon our previously published DeepRescore tool that uses deep learning to improve peptide identification in immunopeptidomics^23^, we present in this paper DeepRescore2, a computational workflow that leverages deep learning-based fragment ion intensity prediction and RT prediction for phosphopeptides to enhance phosphosite localization and PSM rescoring. We benchmark DeepRescore2 against existing state-of-the-art workflows for phosphosite localization and PSM rescoring on a synthetic phosphopeptide dataset. We also demonstrate its application to three real-world biological datasets to improve the sensitivity of phosphopeptide identification, reduce the number of missing values, and boost phosphoproteomics-based biological discoveries.

## Results

### DeepRescore2 workflow

Figure 1 depicts the overall DeepRescore2 workflow, which processes the results of database searching in four steps to improve phosphopeptide identification and phosphosite localization (**Methods**). First, based on the confidently identified PSMs from database searching, RT and fragment ion intensity prediction models are trained using AutoRT^21^,27 and pDeep3^28^, respectively, and then used to predict RTs and MS/MS spectra for all identified peptide sequences with all possible phosphosite localizations (i.e., peptide isoforms). Second, for each peptide isoform, a probability score is computed taking into consideration the PhosphoRS score^16^, RT difference between predicted and experimentally observed RTs, and spectrum similarity between predicted and experimentally observed spectra, and then phosphosite localization is determined based on the combined probability score. Third, PSM rescoring is performed using the semi-supervised Percolator algorithm^29^, which integrates search engine specific features, search engine independent features, and the two deep learning-derived features to improve the accuracy and sensitivity of phosphopeptide identification. Finally, identified PSMs can be manually validated using the visualization tool PDV^30^.

**Figure 1:**
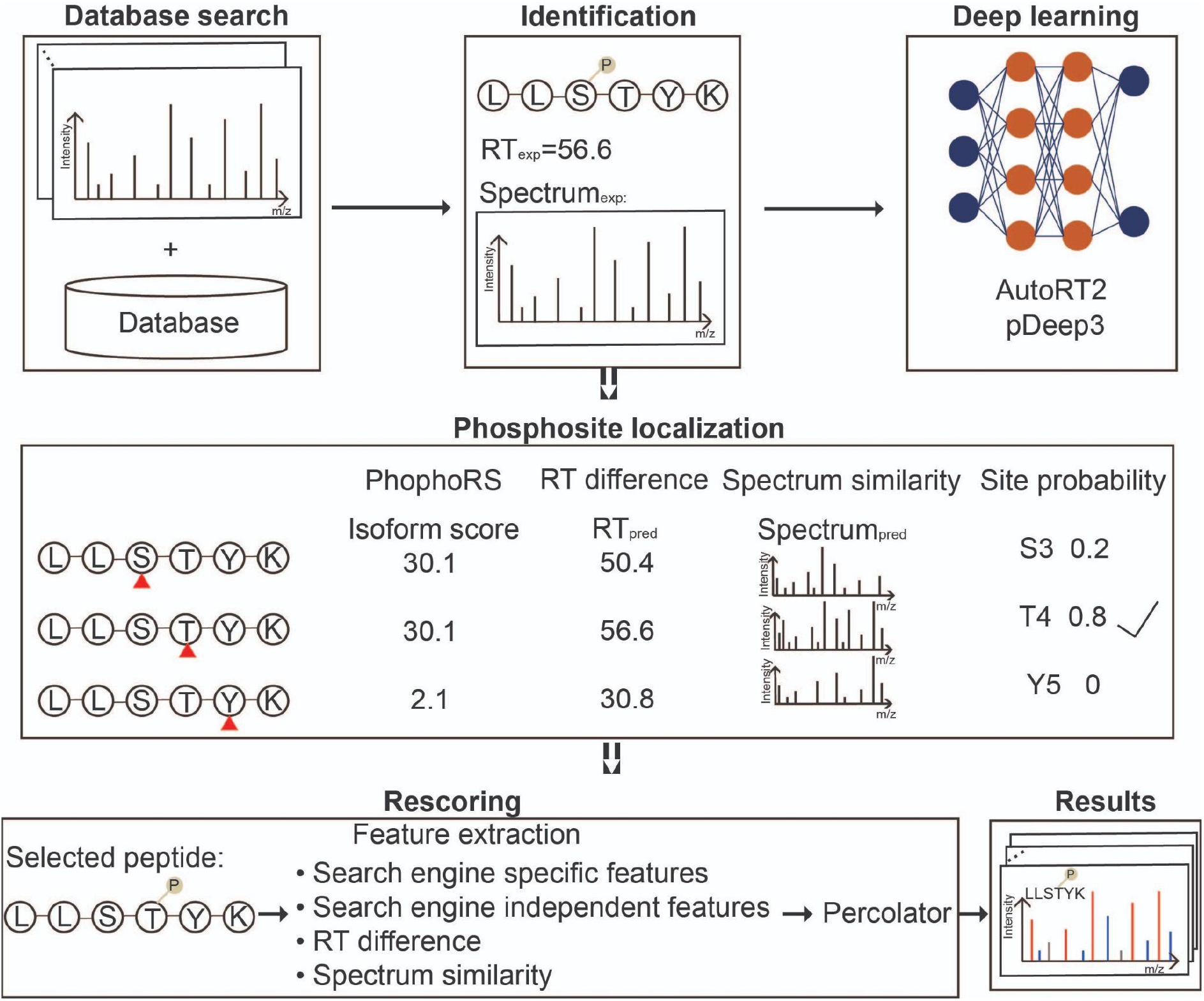
DeepRescore2 workflow for phosphopeptide identification and phosphosite localization. After searching the MS/MS spectra against a protein database using a search engine, DeepRescore2 processes the search results in four steps: 1) RT and fragment ion intensity prediction for all identified peptide sequences with all possible phosphosite localizations using deep learning models fine-tuned with confidently identified PSMs from the database search. 2) Phosphosite localization by combining site localization score from PhosphoRS and deep learning prediction derived spectrum similarity and retention time (RT) difference scores. 3) PSM rescoring using Percolator based on search engine specific features, search engine independent features, and spectrum similarity and retention time difference scores. 4) Visualization of the PSMs by PDV.

### Benchmarking of deep learning-facilitated phosphosite localization

We benchmarked the performance of four site localization methods, PhosphoRS alone (Method 1) and its combination with either spectrum similarity (Method 2), RT difference (Method 3) or both (Method 4), on MaxQuant search results from a synthetic phosphopeptide dataset^31^, in which the ground truth is known and the false localization rate (FLR) can be determined precisely (**Figure 2A, Table S1**). Spectrum similarity was computed based on entropy distance (Entropy)^32^, which outperformed the other five spectrum similarity computation methods including dot product (DP), square root dot product (srDP), spectral contrast angle (SA), Pearson correlation coefficient (PCC), and unweighted entropy distance (unwEntropy) in our comparative analysis (**Methods, Figure S2A**). RT difference was computed based on RT ratio (RTR), which outperformed the alternative method delta RT (DRT) in our comparative analysis (**Methods, Figure S2B**).

**Figure 2:**
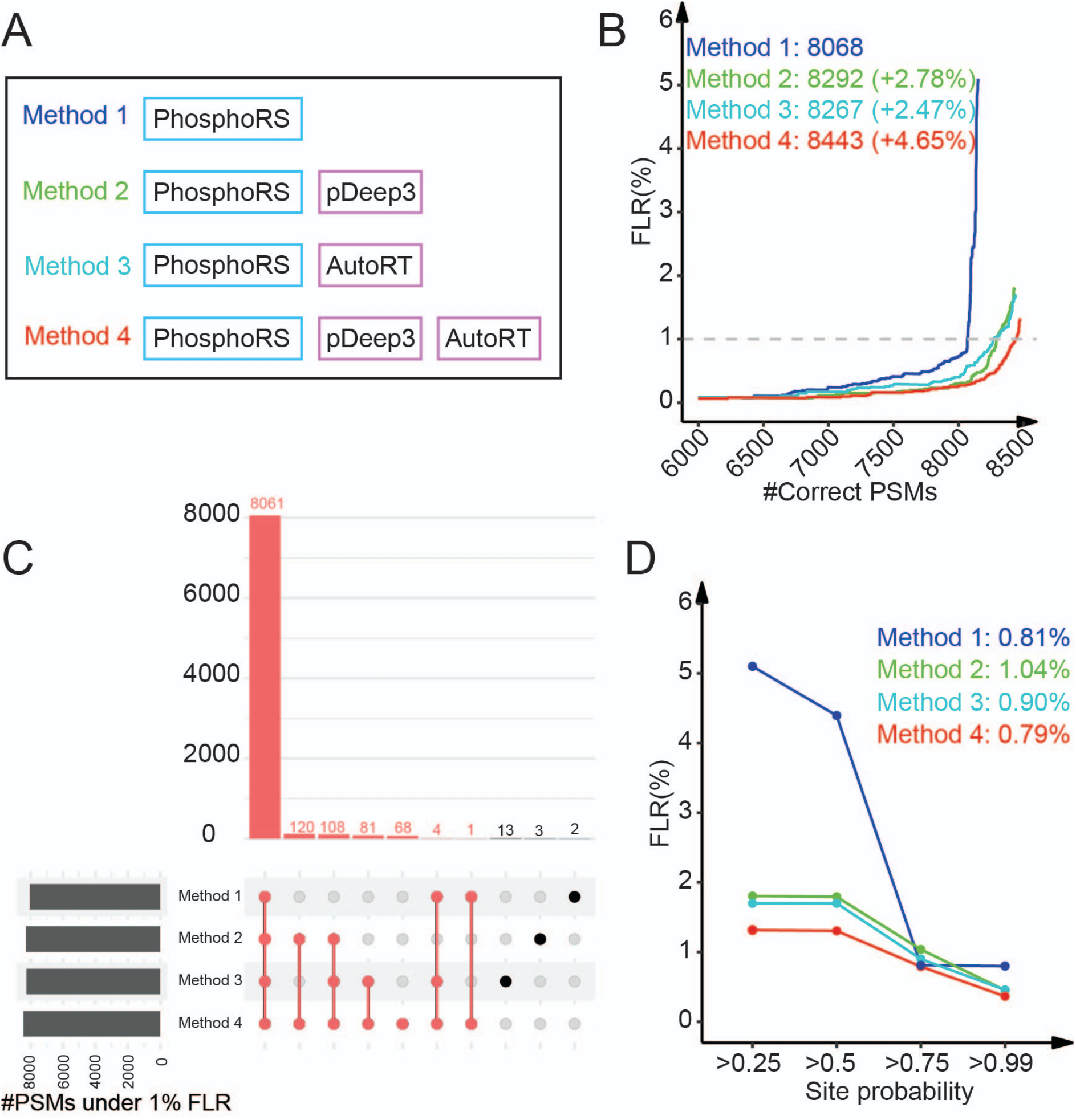
Benchmarking of deep learning-facilitated phosphosite localization on the synthetic dataset. (A) Four site localization methods were benchmarked. (B) The number of correctly localized PSMs at different levels of PSM FLR are shown for the four methods, respectively. The numbers of correctly localized PSMs at 1% FLR and the percent increase compared with Method 1 (phosphoRS) are indicated. (C) UpSet plot comparing the PSM identifications from the four methods. The number of correctly localized PSMs by Method 4 are marked as red. (D) The FLRs at different site probability cutoffs are shown. The FLRs at 0.75 site probability are indicated.

Among the four site localization methods, the ones incorporating deep learning derived features consistently outperformed the PhosphoRS alone method at different levels of FLRs, with the best performance observed for the method simultaneously incorporating both features (**Figure 2B**). At 1% FLR, compared with Method 1, Methods 2, 3, and 4 increased the number of correctly identified PSMs by 2.78%, 2.47%, and 4.65%, respectively. Almost all PSMs correctly identified by Method 1, 2, or 3 were also correctly identified by Method 4 (**Figure 2C**). Moreover, 68 PSMs were correctly identified only by Method 4. These results suggest that Method 4 is the most effective method for phosphosite localization and that complementary information from PhosphoRS score, spectrum similarity, and RT difference are all required to derive the correct phosphosite localization for some PSMs.

We further investigated the relationship between computed site localization probabilities and the true FLR. Among the four methods, Method 4 was associated with the lowest FLRs for all four site localization probability cutoffs investigated, with an FLR of 0.79% observed for the cutoff of 0.75 (**Figure 2D**). Moreover, with the site probability cutoff of 0.75, all other methods were also able to achieve an FLR of around 1%, which is acceptable for typical phosphoproteomics studies. Thus, these results provide empirical evidence to support the use of 0.75 as the site localization probability cutoff in the analysis of real datasets, in which the real FLR is unknown.

### Benchmarking of deep learning-facilitated phosphopeptide identification

Based on MaxQuant search results from the synthetic phosphopeptide dataset, we further benchmarked three phosphopeptide identification methods with different combinations of the site localization and PSM rescoring algorithms, using Method 1 (PhosphoRS only) as the baseline method for comparison (**Figure 3A**). The three methods included PhosphoRS followed by a traditional PSM rescoring algorithm without using deep learning derived features (Method 5), PhosphoRS followed by our proposed DeepRescore algorithm for PSM rescoring (Method 6), and Method 4 followed by DeepRescore (Method 7) (**Table S2**). In DeepRescore, the two deep learning derived features, spectrum similarity and RT difference, were computed based on entropy distance and RT ratio, respectively, based on their superior performance as compared to alternative methods (**Methods, Figure S3A-B**).

**Figure 3:**
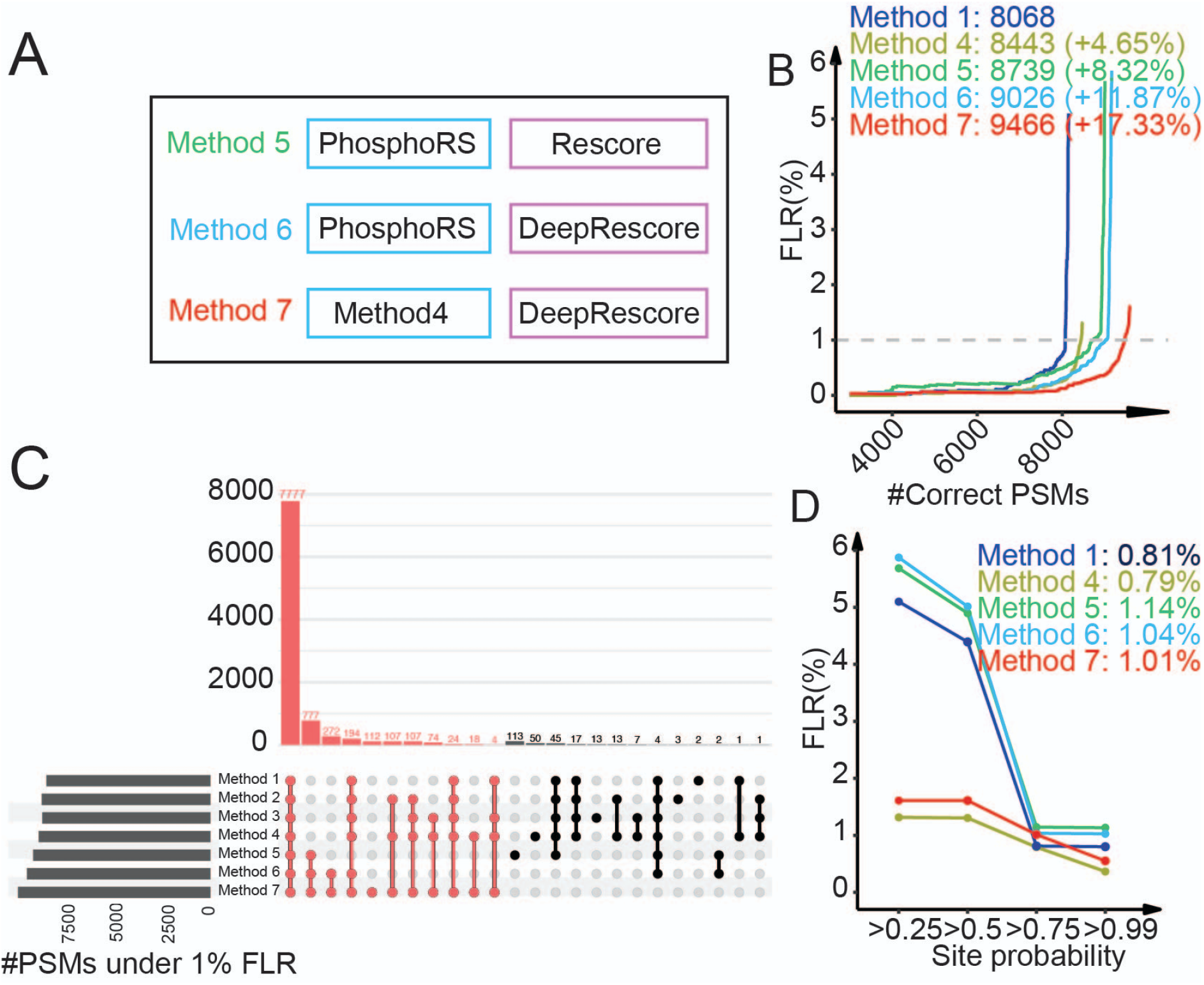
Benchmarking of deep learning-facilitated phosphopeptide identification on the synthetic dataset. (A) Three phosphopeptide identification methods were benchmarked. (B) The number of correctly localized PSMs at different levels of PSM FLR are shown for the three methods, respectively, together with Methods 1 and 4. The numbers of correctly localized PSMs at 1% FLR and the percent increase compared with Method 1 (phosphoRS) are indicated. (C) UpSet plot comparing the PSM identifications from all seven methods. The number of correctly localized PSMs by Method 7 is marked as red. (D) The FLRs at different site probability cutoffs are shown. The FLRs at 0.75 site probability are indicated.

Method 6 and 7 consistently outperformed Method 1 and 5 at different levels of FLRs, and Method 5 outperformed Method 1, but only in the region where FLR was higher than 0.4% (**Figure 3B**). At 1% FLR, compared with Method 1, Methods 5, 6, and 7 increased the number of correctly identified PSMs by 8.32%, 11.87%, and 17.33%, respectively. These increases were much higher than those observed for Methods 2, 3, and 4, suggesting a dominant role of PSM rescoring, especially DeepRescore, in the observed performance gain. The largest performance gain was from Method 7, which leverages deep learning-derived features in both site localization and PSM rescoring. Almost all PSMs correctly identified by the other six methods were also correctly identified by Method 7, and 112 PSMs were correctly identified only by Method 7 (**Figure 3C**). Similar to Methods 1-4, Methods 5-7 were also able to achieve an FLR of around 1% with the site probability cutoff of 0.75 (**Figure 3D**), further supporting the use of 0.75 as the site localization probability cutoff. Together, data from our benchmarking study on a synthetic dataset clearly revealed superior performance for Method 7 compared with other competing methods. Method 7, which leverages deep learning-based retention time and fragment ion intensity predictions in both phosphosite localization and PSM rescoring, was referred to as DeepRescore2 and used in all subsequent studies in this paper.

### Sensitivity improvement in real-world biological datasets

Next, we assessed the performance of DeepRescore2 in real-world biological datasets. We applied it to a label-free dataset from human osteosarcoma cell line U2OS,^33^ and a tandem mass tag (TMT) dataset on uterine corpus endometrial carcinoma (UCEC) from the National Cancer Institute’s Clinical Proteomic Tumor Analysis Consortium (CPTAC) ^34^ (**Methods, Table S1**). To save computational resources, method evaluation was based on three raw files from the U2OS study and one TMT plex from the UCEC study. Both datasets were searched using four search engines Comet, MaxQuant, MSGF+, and X!Tandem, respectively, and the search results were processed using DeepRescore2 or PhosphoRS. Only identifications within the 1% FDR limit at both PSM and phosphopeptide levels and a site localization probability greater than 0.75 were considered as confident identifications.

In the U2OS label-free dataset, when used in combination with DeepRescore2, Comet, MaxQuant, MSGF+, and X!Tandem identified 9,135, 10,380, 8,613 and 8,897 phosphorylated peptides, respectively, which were 31%, 19%, 27%, and 32% higher than the numbers reported when they were used in combination with PhosphoRS (**Figure 4A, Table S3**). In the UCEC TMT dataset, when used in combination with DeepRescore2, Comet, MaxQuant, MSGF+, and X!Tandem identified 27,881, 28,658, 29,082, 25,679 phosphorylated peptides, respectively, which were 40%, 28%, 43%, 46% higher than the numbers reported when they were used in combination with PhosphoRS (**Figure 4B, Table S3**). In both the label-free and the TMT datasets, DeepRescore2 identifications covered almost all PhosphoRS identifications for all search engines (**Figure 4**). All above phosphopeptide-level observations were similarly observed at the PSM level (**Figure S4, Table S3**). Together, these results clearly demonstrate that DeepRescore2 increases the sensitivity of phosphopeptide identification in real-world biological datasets.

**Figure 4:**
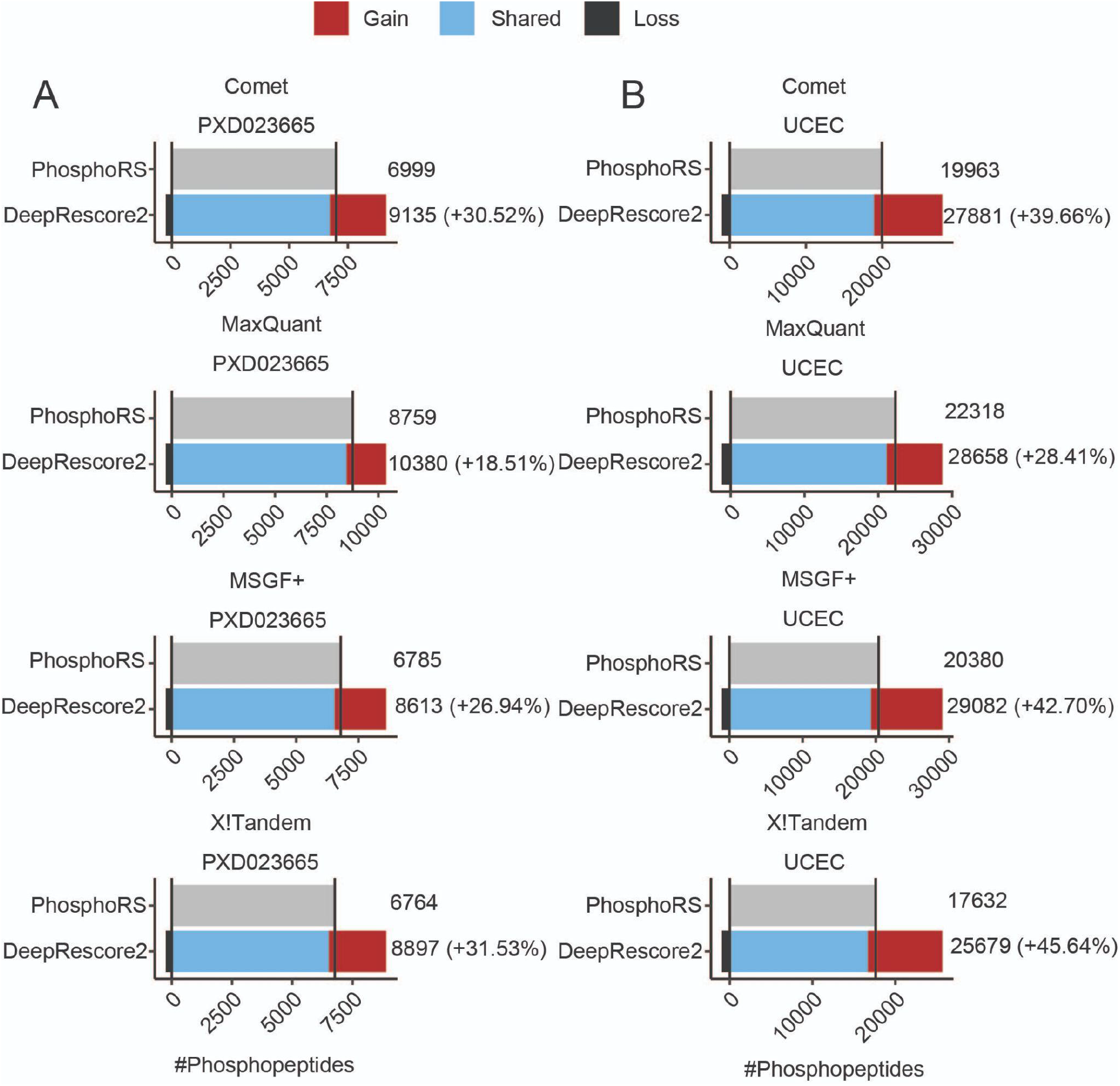
Performance evaluation based on peptide identification in two biological datasets using different search engines in combination with PhosphoRS or DeepRescore2. (A) The numbers of phosphopeptides identified from a label-free phosphoproteomic dataset, PXD023665, by four search engines in combination with phosphoRS or DeepRescore2, respectively. (B) The numbers of phosphopeptides identified from the UCEC TMT phosphoproteomic dataset by four search engines in combination with phosphoRS or DeepRescore2, respectively. Gain: phosphopeptides identified by DeepRescore2 but not PhosphoRS. Shared: phosphopeptides identified by both DeepRescore2 and PhosphoRS. Loss: phosphopeptides identified by PhosphoRS but not DeepRescore2.

### Increased identification of liver cancer-associated phosphosites

Improved phosphopeptide identification is expected to enhance biological discoveries. To demonstrate this potential, we applied DeepRescore2 to a previously published hepatocellular carcinoma (HCC) study^35^, which included TMT-based phosphoproteomic analysis of paired treatment naïve tumors and normal adjacent tissues (NATs) from 159 HCC patients (**Table S1**). The dataset was searched using MaxQuant (**Methods**) and then processed using either PhosphoRS or DeepRescore2. Identifications within the 1% FDR limit at both PSM and phosphopeptide levels and a site localization probability greater than 0.75 were considered as confident identifications. Phosphosite level quantification was performed for confident identifications from PhosphoRS and DeepRescore2 (**Methods**).

Across the whole cohort, DeepRescore2 identified 2,457,685 PSMs and 93,759 phosphopeptides, which increased the PhosphoRS-based PSM and phosphopeptide identifications by 41% and 23%, respectively (**Figure 5A-B**). Importantly, the vast majority of PSM and phosphopeptide identifications reported by PhosphoRS were covered by DeepRescore2 identifications. Phosphosite quantification matrices generated based on DeepRescore2 included more “quantifiable” phosphosites than those based on PhosphoRS identifications across a wide range of non-missing value cutoffs (**Figure 5C**). For downstream analyses, we defined quantifiable sites as those quantified in at least 100 tumor samples and 100 NAT samples. With this definition, DeepRescore2 identified 22,033 quantifiable sites, which was 34% more than those identified by PhosphoRS (**Figure 5D, Table S4**). Only 443 out of the 16,451 (2.7%) quantifiable sites identified by PhosphoRS were not identified by DeepRescore2.

**Figure 5:**
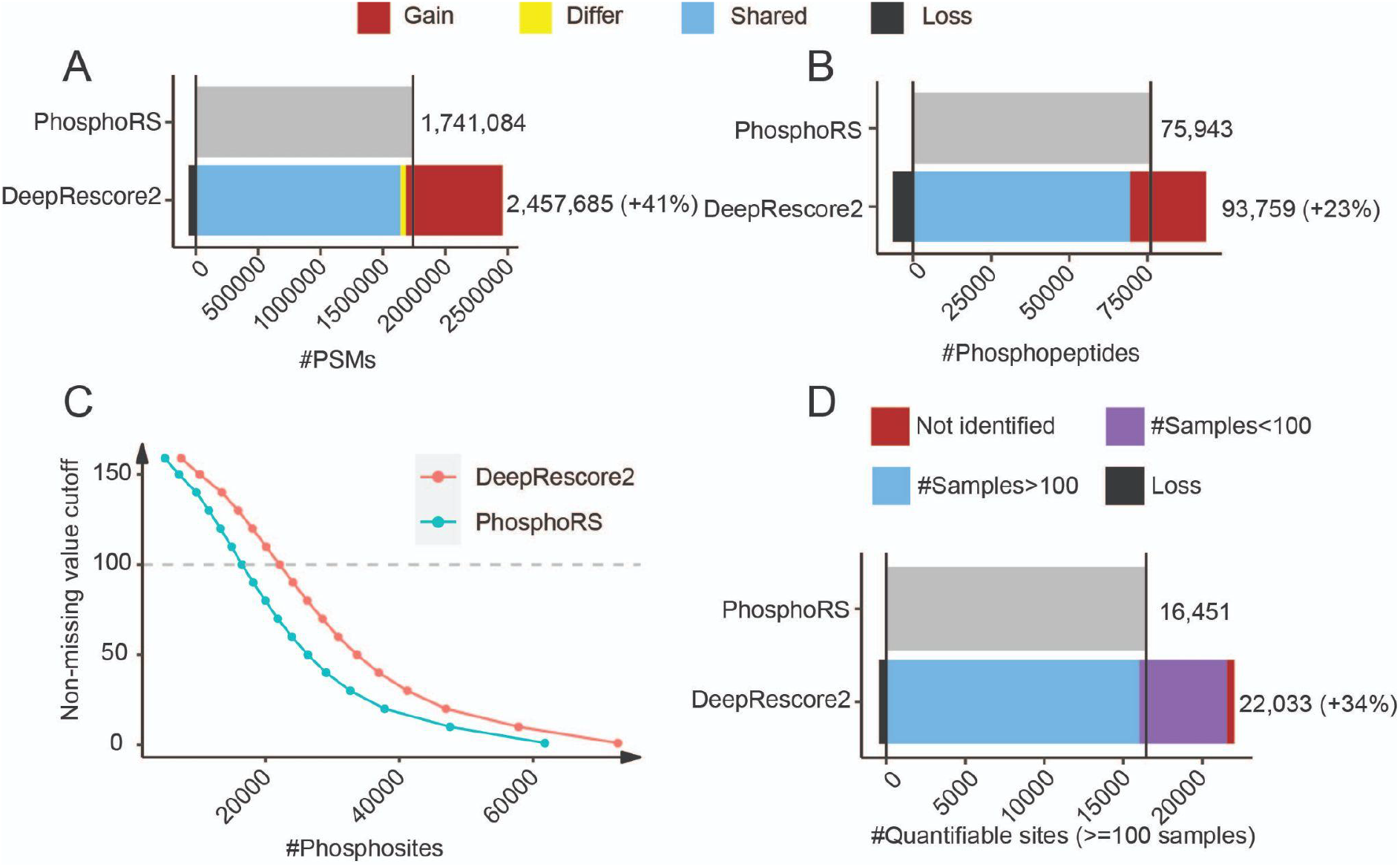
Performance evaluation on a large liver cancer dataset. (A-B) The numbers PSMs (A) and phosphopeptides (B) identified by MaxQuant in combination with phosphoRS or DeepRescore2, respectively. Gain: identified by DeepRescore2 but not PhosphoRS. Shared: identified by both DeepRescore2 and PhosphoRS. Loss: identified by PhosphoRS but not DeepRescore2. Differ: different sites identified by DeepRescore2 and PhosphoRS, respectively. (C) The numbers of “quantifiable” phosphosites based on different levels of non-missing value cutoff, i.e., numbers of samples with a non-missing value in tumors and NATs, respectively. (D) The numbers of quantifiable phosphosites (i.e., quantified in at least 100 tumor samples and 100 NAT samples) in the data tables generated by DeepRescore2 and PhosphoRS, respectively.

To identify liver cancer associated phosphosites, we assessed the differences in phosphosite abundance between tumors and paired NATs. Among the 22,033 quantifiable phosphorylation sites reported by DeepRescore2, 5,021 were significantly increased and 4,929 were significantly decreased in tumors compared to paired NATs (adj. p < 0.01, Wilcoxon signed-rank test). Of these, only 3,567 (71%) and 3,498 (71%) were also found to be significantly altered based on PhosphoRS-derived data, and others were not identified (95+113), not quantifiable (967+1,097), or not significantly altered (392+221) (**Figure 6A, Table S4**). Significantly altered phosphosites that were not identified or not quantifiable in PhosphoRS-derived data included known functional sites on well-established cancer genes such as CHEK2_S379, which induces the enzymatic activity of CHEK2^36^, and EGFR_S1071, which drives EGFR desensitization^37^, as well as sites on less well-studied genes such as RIPK2_S176, which induces the enzymatic activity of RIPK2^38^ (**Figure 6B**).

**Figure 6:**
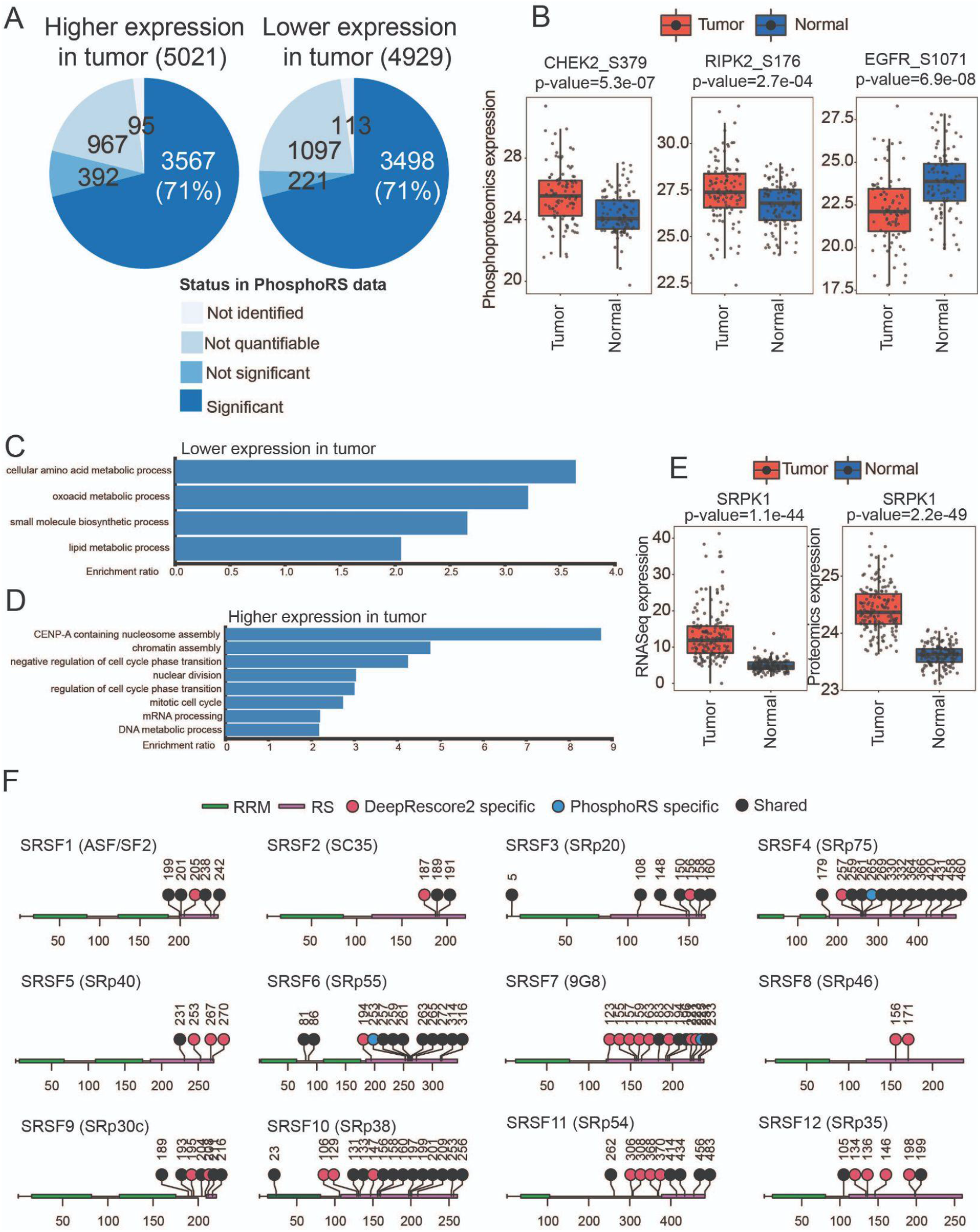
Differentially expressed phosphosites between tumor and NAT samples in the liver cancer dataset. (A) The numbers of significantly altered (FDR<0.01) phosphosites identified in DeepRescore2-derived data and their status in the analysis of PhosphoRS-derived data. (B) Comparison of phosphosite abundance in tumors and NATs for CHEK2_S379, RIPK2_S176, and EGFR_S1071, respectively. (C-D) Gene Ontology enrichment analysis results of the parent proteins of the 5,021 significantly down-regulated phosphosites (C) and those of the 4,929 significantly up-regulated phosphosites (D). (E) Comparison of SRPK1 abundance between tumors and NATs at mRNA and protein levels, respectively. (F) Differentially expressed phosphosites detected in DeepRescore2- or PhosphoRS-derived data on 12 SR proteins. *p* values are based on Wilcoxon rank sum test.

At the pathway level, parent genes of the 4,929 decreased sites were enriched in different metabolic processes (**Figure 6C**), consistent with the essential role of liver in metabolism. Parent genes of the 5,021 increased sites were enriched in biological processes related to CENP-A containing nucleosome assembly, chromatin assembly, negative regulation of cell cycle phase transition, nuclear division, regulation of cell cycle phase transition, mitotic cell cycle, mRNA processing, DNA metabolic process, etc (**Figure 6D**). Protein phosphorylation plays an important role in many mRNA processing events, including pre-mRNA splicing^39^. Phosphorylation status greatly modulates the activity of SR proteins, a family of nuclear RNA binding proteins involved in the regulation of both constitutive and alternative splicing^40^. In our tumor vs NAT comparison, 98 phosphosites from the 12 SR proteins were significantly increased in tumors. Moreover, 78 (80%) of the 98 phosphosites were on serines located in the RS domain. This region is specifically recognized by SRPK1^41^, the most well-studied SR protein kinase^42^. In accordance with this observation, SRPK1 showed significantly increased mRNA and protein abundance in tumors compared with NATs in this cohort (**Figure 6E**). Thus, data from both kinase and substrates support a role of SPRK1 activity in liver cancer development. Importantly, only 55 out of the 78 phosphosites (71%) in the RS domain were identified as significantly increased in the analysis of the PhosphoRS-derived dataset (**Figure 6F**). Together, DeepRescore2 provides a more comprehensive catalog of liver cancer associated phosphosites, linking functional phosphosites and biological processes to liver cancer oncogenesis, and expanding the list of putative substrates of liver cancer-associated SRPK1 activity.

### Increased identification of prognosis-associated phosphosites in liver cancer

To identify prognosis-associated phosphosites in liver cancer, we further performed survival analysis based on overall survival (OS) data of the 159 HCC patients. Among the 22,033 quantifiable phosphorylation sites reported by DeepRescore2, 420 were significantly associated with poor prognosis (p<0.01, hazard ratio [HR] >2, log rank test) and 202 were significantly associated with good prognosis (p<0.01, HR<0.5). Of these, only 176 (42%) and 79 (39%) were also found to be significantly associated with prognosis based on PhosphoRS-derived data, and others were not identified (9+3), not quantifiable (95+38), or not significantly associated (140+82) (**Figure 7A, Table S4**). Among the top 10 poor prognosis-associated phosphosites with the largest HRs, only seven showed significant association with OS in PhosphoRS-derived data (**Figure 7B**). Moreover, eight out of the 10 phosphosites had a larger hazard ratio than those computed based on cognate mRNA and protein measurements (**Figure 7B**). In particular, NAV3_S1190 and MIEF1_S79 were significantly associated with poor prognosis based on DeepRescore2-but not PhosphoRS-derived data, while their cognate mRNA and protein were not significantly associated with OS (**Figure 7C-D**), suggesting a specific contribution of phosphorylation to the observed survival associations.

**Figure 7:**
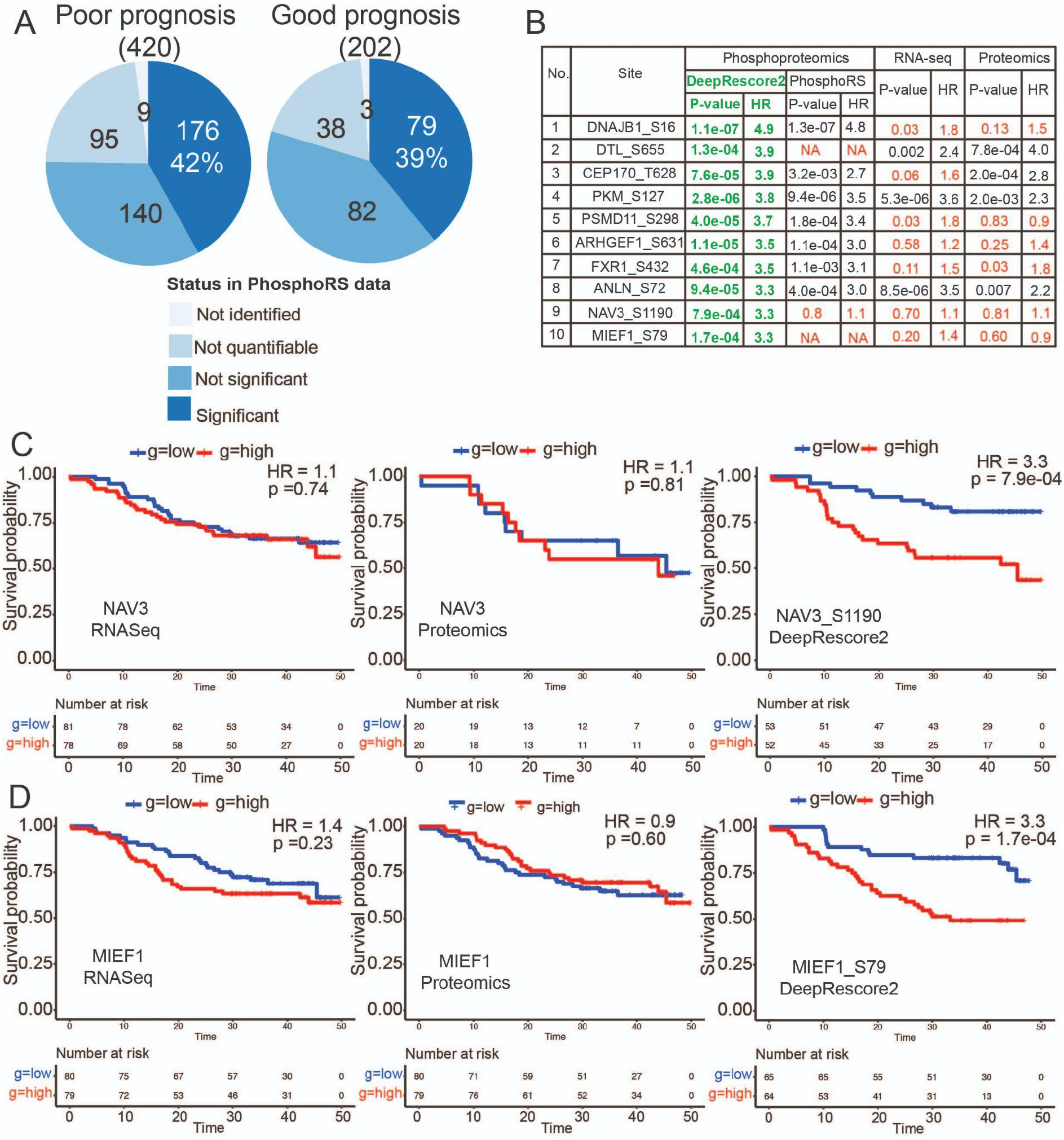
Prognosis-associated phosphosites in the liver cancer dataset. (A) The numbers of phosphosites significantly associated with prognosis (p<0.01 and HR>2 [poor prognosis] or HR<0.5 [good prognosis]) in DeepRescore2-derived data and their status in the analysis of PhosphoRS-derived data. (B) Survival data for the top 10 poor prognosis-associated phosphosites in DeepRescore2-derived data and results for their parent genes based on RNA and protein measurements, respectively. The green color indicates the DeepRescore2 results. The red color indicates that the site is not identified, not quantified, or not significant. (C) Kaplan Meier curves for patients stratified based on the median abundance of NAV3 RNA, NAV3 protein, and NAV3_S1190 phosphorylation, respectively. (D) Kaplan Meier curves for patients stratified based on the median abundance of MIEF1 RNA, MIEF1 protein, and MIEF1_S79 phosphorylation, respectively. *p* values are based on the log-rank test.

### EGFR hyperactivation as a new target in poor-prognosis liver cancer

Since kinases are important therapeutic targets, we next performed kinase activity inference for the 159 HCC tumor samples based on DeepRescore2- and PhosphoRS-derived phosphoproteomics datasets, respectively (**Methods**). Kinase activity was quantified for 134 and 120 kinases/kinase families based on DeepRescore2- and PhosphoRS-derived data, respectively. To identify prognosis-associated kinases, we performed survival analysis using kinase activity scores derived from DeepRescore2 and PhosphoRS data, respectively.

The activities of eight kinases (EGFR, CDK7, AMPK-family, CDK6, CDC7, CDK2, ATR, and RPS6KA5) were significantly associated with OS (p<0.05, HR>2 or <0.5, log rank test) in one or both analyses (**Table S5**), and analysis results based on the two datasets were comparable for the vast majority of kinases (**Figure 8A**). One notable outlier was EGFR, for which higher inferred activity was associated with shorter OS, but only when the inference was made based on DeepRescore2 data (**Figure 8A-C**).

**Figure 8:**
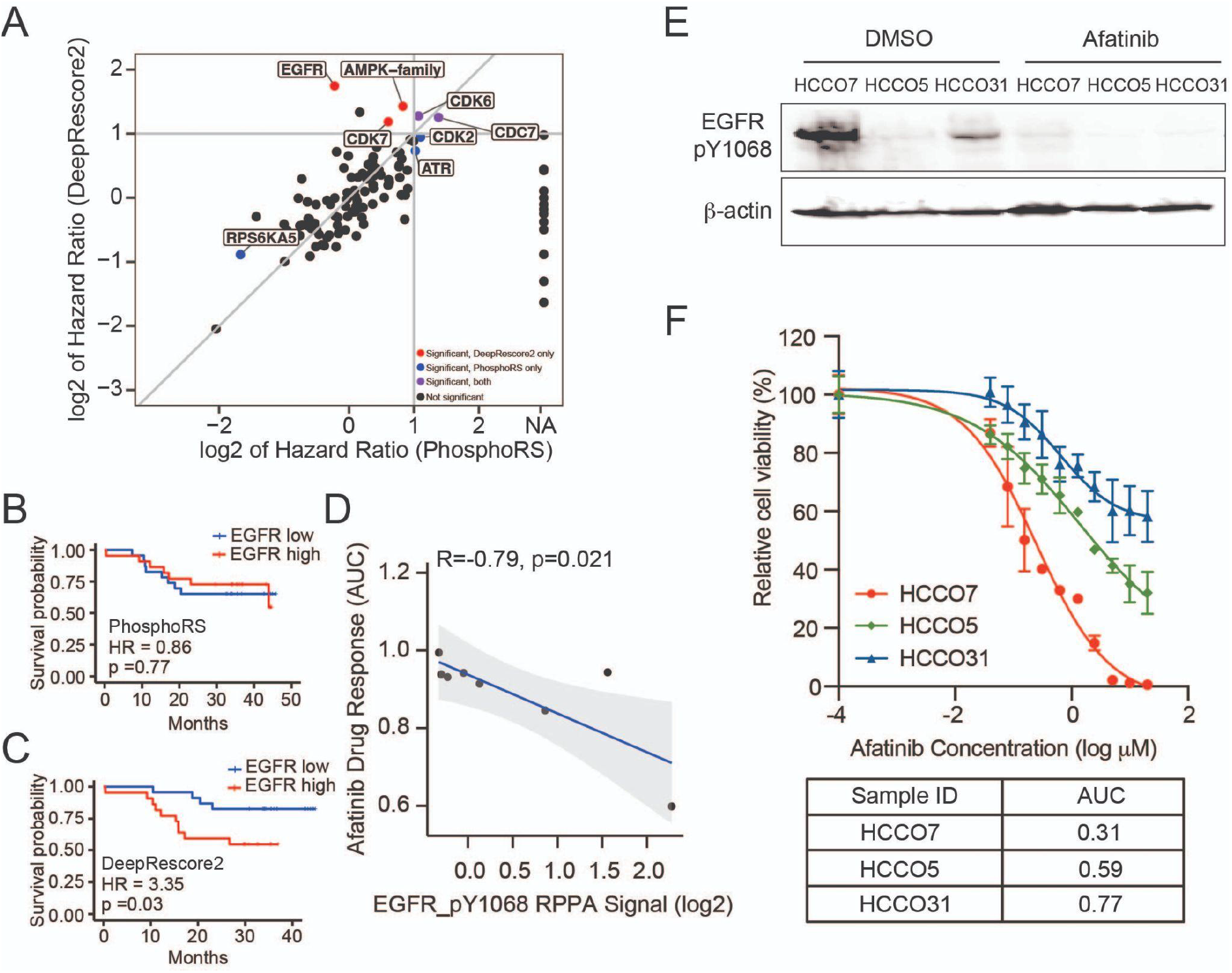
Kinase activity analysis identifies EGFR hyperactivation as a new target in poor-prognosis liver cancer. (A) Scatterplot comparing hazard ratios of kinase activities inferred based on DeepRescore2- and PhosphoRS-derived data, respectively. (B) Kaplan Meier curves for patients stratified based on median abundance of EGFR activity inferred based on PhosphoRS-derived data. (C) Kaplan Meier curves for patients stratified based on median abundance of EGFR activity inferred based on DeepRescore2-derived data. (D) Correlation between afatinib response quantified based on AUC and EGFR_Y1068 abundance measured by RPPA. (E) Western blot analysis of EGFR_Y1068 on three HCC organoids, HCCO7, HCCO5, HCCO31, treated with DMSO or 5µM afatinib for 2 hours. (F) Relative cell viability as a function of afatinib concentration in the three organoids. Data were from three replicates.

To examine whether EGFR hyperactivation could serve as a target for anti-cancer interventions in HCC, liver cancer cell line data from DepMap was used to correlate Reverse Phase Protein Array (RPPA) measurement of EGFR_Y1068, a site that is crucial to EGF-induced Ras/MAPK signaling^43^, with response to afatinib, an EGFR-specific tyrosine kinase inhibitor. We observed a strong negative correlation between EGFR_Y1068 abundance and afatinib response as quantified by the area under the curve (AUC, smaller AUC corresponds to higher sensitivity) across the eight liver cancer cell lines (*R* =− 0. 79, *p* = 0. 021, spearman correlation, **Figure 8D)**, suggesting that EGFR_Y1068 levels could be a potential marker of response to EGFR inhibition.

To further confirm this finding from cell line-based high-throughput study, we identified three HCC organoids with varied levels of EGFR_Y1068 phosphorylation by Western blot (**Methods**). HCCO7 expressed a high level of EGFR_Y1068, whereas HCCO31 and HCCO5 had much lower levels of EGFR_Y1068 abundance (**Figure 8E**). Upon afatinib treatment, a detectable decrease of EGFR_Y1068 was observed in all models, confirming drug target engagement (**Figure 8E**). HCCO7 demonstrated high sensitivity to afatinib (AUC = 0.31, **Methods**), while the other organoids were less sensitive (**Figure 8F**). These results corroborate the results from the DepMap cell line analysis and support the use of EGFR_Y1068 as a marker of response to EGFR inhibitors in liver cancer. Together, our data demonstrate the power of improved phosphosite identification in improving kinase activity inference, leading to the identification of poor-prognosis-associated hyperactivated kinases as new targets for drug repurposing or development.

## Discussion

We developed DeepRescore2, a computational workflow that leverages deep learning prediction to improve phosphopeptide identification and phosphosite localization in phosphoproteomics data analysis. DeepRescore2 substantially increases the sensitivity of phosphopeptide identification in both synthetic and real-world biological datasets, and improved identification leads to an in-depth understanding of kinase signaling and deeper biological discoveries.

A key strength of DeepRescore2 is the integrative workflow that leverages complementary computational techniques including PhosphoRS scoring, semi-supervised machine learning through Percolator, and deep learning-based prediction of phosphopeptide RT and fragment ion intensities. Based on a synthetic dataset with known ground truth, our benchmarking study clearly showed that the combination of all these techniques collectively contributed to the ultimate best performance. Performance gain was observed when incorporating deep learning prediction in either phosphosite localization or PSM rescoring, but the latter provided a stronger sensitivity increase.

During the benchmarking process, we also comprehensively assessed the impact of different methods for computing spectrum similarity and RT difference on the performance of site localization and PSM rescoring (**Figure S2-S3**). Our analyses revealed a substantial impact of method selection on the performance of both tasks. For spectrum similarity computation, entropy distance, a similarity scoring method recently proposed for metabolomics data analysis^32^, showed the best performance in our evaluation. For RT difference computation, a new method, RT ratio, outperformed the routinely used delta RT method. Although not the focus of this study, these results should be of particular interest to the proteomics and mass spectrometry community.

Applying DeepRescore2 to three real-world biological datasets demonstrated its application to both label-free and TMT phosphoproteomics datasets, and its flexibility to be combined with different search engines. Importantly, improved sensitivity in phosphopeptide identification directly translated into increased biological discoveries. Most of the phosphosites significantly associated with liver cancer development and prognosis have very limited information in the literature, opening new opportunities for further investigation. Increased phosphosite identification also led to improved kinase activity inference. One important finding is that EGFR hyperactivation is not only associated with poor-prognosis in liver cancer, but also associated with increased sensitivity to afatinib in both cell lines and organoids. Afatinib is approved for non-small cell lung cancer harboring EGFR mutations. It is also being investigated as a tissue-agnostic drug in a biomarker guided phase 2 basket clinical trial where patients with tumors, regardless of location, harboring EGFR activating mutations are matched to treatment with afatinib (NCT02465060). Although EGFR inhibitors have not been approved for liver cancer, our finding may accelerate mechanism-based drug repurposing by incorporating EGFR_Y1068 as a patient stratification marker for clinical investigation.

One limitation of the study is that our new method was combined only with the phosphosite localization algorithm phosphoRS. We chose phosphoRS as the baseline method because it showed relatively higher sensitivity compared with other phosphosite localization algorithms in previous evaluation studies^26,31,44^. During our manuscript preparation, a new phosphosite localization algorithm, AScorePro, has been published and showed superior performance compared with its predecessor AScore^45^. Because the deep learning-derived features are largely independent of the information used in AscorePro, we expect that combining deep learning-derived features with AscorePro could also improve phosphosite localization. However, this will need to be formally tested in the future. It would also be interesting to leverage the new Iterative Synthetically Phosphorylated Isomers (iSPI) resource^45^ in future benchmarking efforts. Moreover, DeepRescore2 could be further improved by integrating additional deep learning-derived features, such as phosphosite probability prediction^21^. With the increasing use of other PTM profiling in biomedical research, DeepRescore2 can also be expanded to support other PTMs by incorporating deep learning derived features trained on data of other PTMs, which will further broaden the impact of the DeepRescore2 workflow.

## Methods

### Datasets

One liquid chromatography tandem mass spectrometry (LC-MS/MS) dataset of synthetic phosphopeptides and three real-world LC-MS/MS phosphoproteome datasets from biological samples were used in this study. Raw MS data of the synthetic dataset were downloaded from the PRIDE database (https://www.ebi.ac.uk/pride/) with the accession key PXD000138, which included 96 libraries generated from 96 seed peptides^31^. For each seed peptide, the seed position was synthesized with either serine, threonine or tyrosine or their phosphorylated forms, and the amino acids before or after the seed position were permuted with 20 amino acids to generate up to 2,400 different phosphorylated or non-phosphorylated peptides for each library. In this study, we used libraries 1-11, which were generated within a single day (**Table S1**). One of the three biological datasets was a label-free phosphoproteome dataset, and the other two were TMT phosphoproteome datasets. The label-free dataset was from the U2OS cell line. Raw data were downloaded from the PRIDE database with the accession key PXD023665^33^, and three raw files were used in this study for method evaluation (**Table S1**). Raw data of the TMT datasets were downloaded from the Proteomic Data Commons (PDC, https://pdc.cancer.gov/pdc/). The first one is a TMT10-labeled phosphoproteome dataset from the CPTAC UCEC study^34^ with the accession key PDC000126, and the first plex including 12 fractions were used in this study for method evaluation (**Table S1**). The second dataset is a TMT10-labeled phosphoproteome dataset from the International Cancer Proteogenome Consortium (ICPC) HBV-related HCC study^35^ with the accession key PDC000199, and all 33 plexes were used in this study for biological discovery (**Table S1**). All the raw MS/MS data files used were converted to MGF files using ProteoWizard (v3.0.19014).

### Database searching

For the synthetic dataset, we followed the original study^31^ and used MaxQuant (v1.6.5.0) to search the MS/MS data against the human IPI v3.72 database concatenated with the default MaxQuant contaminant database. Oxidation of methionine (+15.9949) and phosphorylation of serine, threonine and tyrosine (+79.9663) were used as variable modifications, and no fixed modification was used. Trypsin/P was used as the digestion enzyme and up to 4 missed cleavages were allowed. The default settings for other parameters were used, but with all statistical filters in MaxQuant, including PSM, protein, and site level FDRs, disabled in order to collect all possible candidate PSMs for downstream analyses.

For the U2OS and UCEC datasets, MS-GF+ (v2019.02.28), Comet (2018.01 rev. 4), X!Tandem (v 2017.2.1.2), and MaxQuant (v1.6.5.0) were used to evaluate the performance of DeepRescore2 in combination with different search engines. For the HCC dataset, MaxQuant (v1.6.5.0) was used as the search engine. These datasets were searched against the human protein database downloaded from Uniprot (v 20190214) concatenated with the default contaminant database in the MaxQuant analysis. Parameters for database searching were set as follows: Oxidation of methionine (+15.9949) and phosphorylation of serine, threonine and tyrosine (+79.9663) were used as variable modifications, and carbamidomethylation of cysteine (+57.02146) was used as fixed modification. For the TMT datasets, TMT labeling (+229.1629) of peptide N-termini and lysine residues were specified as fixed modifications. Trypsin/P was used as the digestion enzyme. Up to 2 missed cleavages were allowed for the U2OS and HCC data, and up to 4 missed cleavages were allowed for the UCEC data. The precursor mass tolerance was set as 20 p.p.m and the MS/MS mass tolerance was set as 0.02 Da.

### Peptide RT and fragment ion intensity prediction

AutoRT^21^,27 was used for RT prediction, and pDeep3^28^ was used for fragment ion intensity prediction for each peptide sequence. Briefly, the phosphorylation-specific base model of AutoRT available at https://github.com/bzhanglab/AutoRT was fine-tuned through transfer learning using experiment-specific data to create experiment-specific models. The phosphorylation-specific base model of pDeep3 available at https://github.com/pFindStudio/pDeep3 was fine-tuned through transfer learning using experiment-specific data to create experiment-specific models. To construct experiment-specific training datasets, we included both phosphorylated and non-phosphorylated peptide identifications passing 1% FDR filtering at both PSM and peptide levels, and the phosphorylated peptide identifications were also required to have higher than 0.75 phosphoRS site localization probability. The experiment-specific models were then used to predict RT and MS/MS spectrum for all peptide isoforms of all identified peptides.

### Deep learning-facilitated site localization

For each phosphorylated site candidate *i* in an identified peptide sequence corresponding to spectrum *S*, two deep learning derived scores were used to adjust the phophosRS localization score of this candidate site *i* (PhosphoRSScore_*i*_). The first deep learning derived score is the spectrum similarity (SS) between a predicted MS/MS spectrum *S*_*p*_ and corresponding experimental MS/MS spectrum *S*_*e*_. Several MS/MS spectrum similarity calculation methods were investigated, including DP, srDP, SA, PCC, unwEntropy^32^, and Entropy^32^. Let *I*_*p*_ and *I*_*e*_ be two vectors containing the peak intensities from *S*_*p*_ and *S*_*e*_, respectively; similarity between *S*_*p*_ and *S*_*e*_ can be calculated by the six different scoring methods as follows:

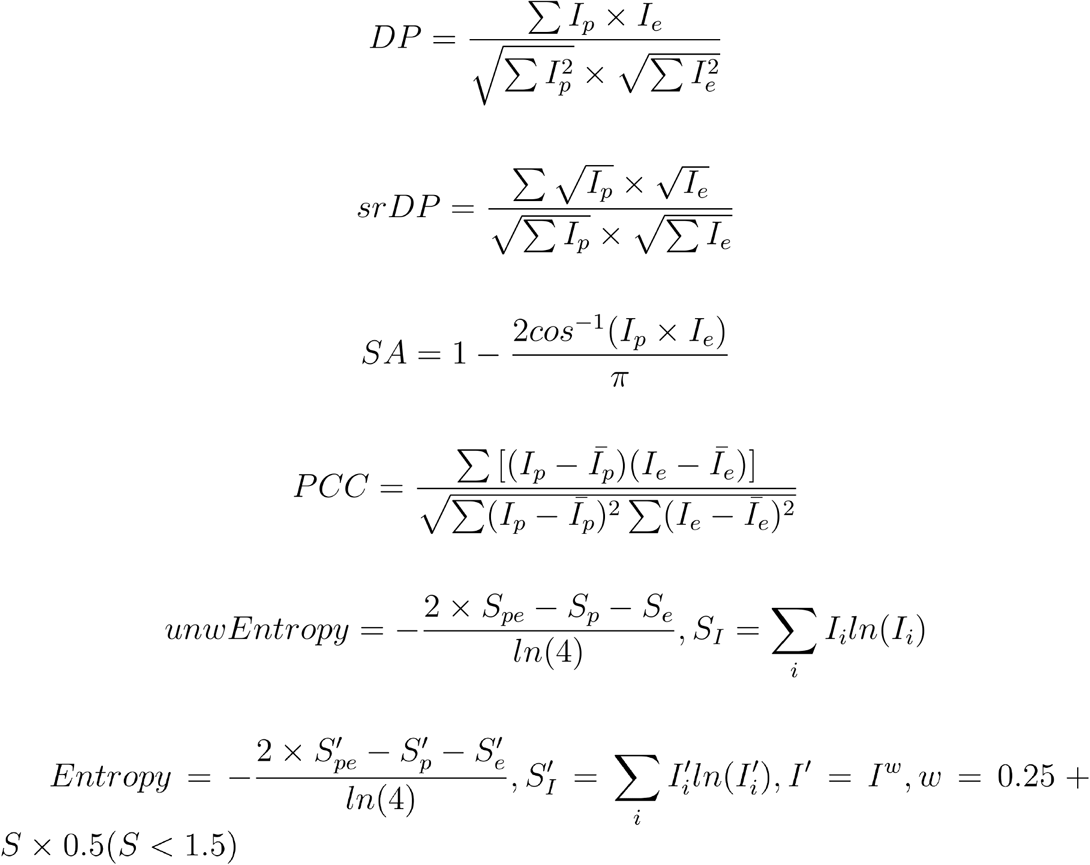

DP, PCC, unwEntropy and Entropy were calculated using the Spectral Entropy python package^32^. srDP and SA were calculated by python scripts following the equations.

The second deep learning derived score is the retention time difference between predicted RT (*RT*_*p*_) of the identified peptide sequence with phosphorylated site candidate *i* and experimentally observed RT (*RT*_*e*_) of the corresponding spectrum. Two methods were used to calculate the difference between *RT*_*p*_ and *RT*_*e*_. The first is delta RT (DRT):

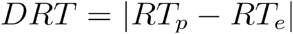

The closer DRT is to 0, the closer *RT*_*p*_ is to *RT*_*e*_. The second is RT ratio (RTR):

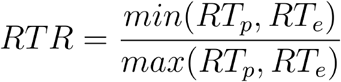

The closer RTR is to 1, the closer *RT*_*p*_ is to *RT*_*e*_.

For a candidate site *i* of an identified peptide sequence corresponding to spectrum *S*, the following equations were used to adjust its PhosphoRS localization score by integrating SS score and DRT score or RTR score :

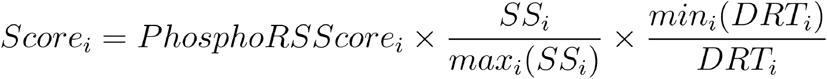

or:

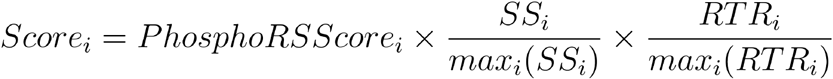

where *max*_*i*_(*SS*_*i*_) and *max*_*i*_(*RTR*_*i*_) are the maximum SS and RTR for all the phosphorylated site candidates in the identified peptide sequence, respectively, and *min*_*i*_(*DRT*_*i*_) is the minimum for all the phosphorylated site candidates in the identified peptide sequence.

Finally, the site localization probability of phosphorylated site candidate *i* was calculated by transforming the localization score through the base 10 SoftMax transformation formula as previously described^26^:

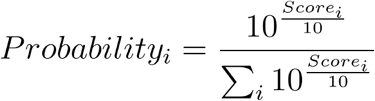

### Deep learning-facilitated PSM rescoring

The phosphorylated site candidate with the largest site localization probability was selected as the site identification of spectrum *S*. Following the DeepRescore strategy^23^, three feature sets (**Table S2**) were extracted from the original PSM identifications or calculated based on the selected phosphorylated site and non-phosphorylated identifications. The first set comprised search engine-specific features, the second set comprised search engine-independent features, and the last set comprised deep learning-derived features, including both RTR and SS computed based on the entropy distance. Using the semi-supervised support vector machine (SVM)-based Percolator rescoring model, all above features were integrated to compute a new score for each PSM.

### False localization rate (FLR) calculation for the synthetic dataset

For each method, all phosphorylated PSMs with correctly identified sequence were sorted by the site localization probability in descending order, and the FLR was calculated as:

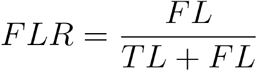

where TL and FL are the numbers of true and false localization sites at the site localization probability cutoff.

### Phosphosite level quantification

The single site-level quantitation procedure was written in Python 3 (Python Software Foundation. Python Language Reference http://www.python.org), along with third-party libraries NumPy^46^ and Pandas (mckinney-proc-scipy-2010, reback2020pandas).

First, any identical modified peptide sequences (different PSMs; different scans with the same modified amino acid sequence) are combined into one record. Quantities are summed, and the best database search score is kept, creating a table unique on modified sequence. Next, the table is expanded to produce a separate record for each modification (e.g., a doubly phosphorylated modified sequence is duplicated into two records, one specific to each position). From here, a 15-mer site identifier is calculated for each record. Final site level quantitation is produced by summing each group of 15mers. This procedure is performed independently for each entry in the reference protein database. Phosphosite abundances were log2 transformed and median normalized for downstream analyses.

### Kinase activity inference

Kinase activity scores were inferred for each data table using the Kinase-Substrate Enrichment Analysis (KSEA) algorithm^47^ implemented in R. Kinase and kinase family target phosphorylation sites collected from several sources and used to evaluate the GPS 5.0 target prediction model were used for the inference here (Table S4 from Wang et al.^48^). For this study, we required measurements for at least five kinase targets in a given sample for the HCC dataset for kinase activity inference.

### Liver cancer organoid culture and afatinib response testing

Liver cancer specimens were collected from patients who underwent surgical resection at Zhongshan Hospital of Fudan University with signed informed consent forms. Liver cancer tissues were minced and digested at 37°C in phosphate buffer saline (PBS, Gibco) containing Collagenase Type IV (5mg/ml, Gibco) for 30-60 min with gentle shaking. After digestion, the suspension was filtered through a 100-mm cell strainer and centrifuged consecutively at 1000 rpm, 800 rpm, and 600 rpm for 5 min, respectively. The pellet was resuspended in cold organoid culture medium and then mixed 1:2 with Matrigel (Corning) to reach a density of 4000 cells per 50 mL before seeding into a 24-well culture plate. After Matrigel solidification at 37°C, organoid culture medium was added to each well and organoids were cultured in a humidified incubator at 37°C with 5% CO_2_.

For Afatinib (Selleck) response testing, organoids were gently digested and seeded into 384-well plates (Corning) at a density of 500 cells per well in a volume of 15 ml (1:1 mixture of culture medium and Matrigel) by Multidrop Combi Reagent Dispenser (Thermo Fisher Scientific). After an incubation at 37°C to make the Matrigel-medium mixture solidify,

35 ml of the pre-warmed culture medium was added into each well. After 72-hour incubation, organoids were treated with serial dilutions of Afatinib using a D300e Digital Dispenser (Tecan). After another 72-hour drug treatment, 20 ml of CellTiter Glo 3D (Promega) was added to each well followed by measuring the luminescent signals using an EnVision Multilabel Reader (PerkinElm) to determine the cell viability. Three replicative wells were measured. AUC was calculated with the equation:

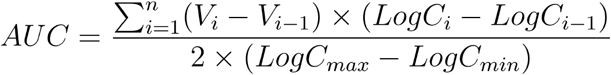

where *C*_*i*_ stands for each concentration and *V*_*i*_ stands for the relative cell viability at *C*_*i*_. *C*_*max*_ and *C*_*min*_ were the maximum and minimum concentration for each drug in our screening.

### Western blot analysis of EGFR_Y1068

For the analysis of phospho-EGFR inhibition, 5 µM afatinib was added into the culture medium, and organoids were collected after incubation for 2 hours. The organoid samples were collected and lysed in RIPA buffer supplemented with protease and phosphatase inhibitor cocktail (Beyotime). The samples were quantified by BCA Protein Assay Kit (Beyotime), separated by 10% SDS-PAGE (Beyotime), and transferred to a PVDF membrane (Millipore). The membrane was blocked with QuickBlock™ Western (Beyotime) and incubated with primary antibody (Phospho-EGF Receptor (Tyr1068) (D7A5), Cell Signaling Technology, CST-3777, 1:1000 dilution) at 4°C overnight followed by secondary antibody incubation. The protein bands were visualized using ChemistarTM ECL Western Blotting Substrate (Tanon).

## Code availability

Source code of DeepRescore2 is available at GitHub (https://github.com/bzhanglab/DeepRescore2).

## Data availability

Phosphoproteomics data used in this study were downloaded from public data repositories as described in the Methods section.

## Acknowledgements

This study was supported by grants U24 CA210954, U24 CA271076, and R01 CA245903 from the National Cancer Institute (NCI), the Cancer Prevention & Research Institutes of Texas (CPRIT) award RR160027, and funding from the McNair Medical Institute at The Robert and Janice McNair Foundation. B.Z. is a CPRIT Scholar in Cancer Research and a McNair Scholar. BCM Mass Spectrometry Proteomics Core is supported by the Dan L. Duncan Comprehensive Cancer Center Award (P30 CA125123) and the CPRIT Core Facility Awards (RP170005 and RP210227).

## Author contributions

B.Z. and B.W. conceived the study. X.Y., B.Z., and B.W. designed the algorithm. X.Y. implemented the algorithm and analyzed the data with the help from B.Z., A.S., E.J. and J.T.L.. S.J. performed the experimental validation under the supervision of Q.G.. B.Z. and X.Y. wrote the manuscript. All authors read and approved the final manuscript.

## Competing interests

The authors declare no competing financial interests.

## Supplemental Figures

**Figure S1:**
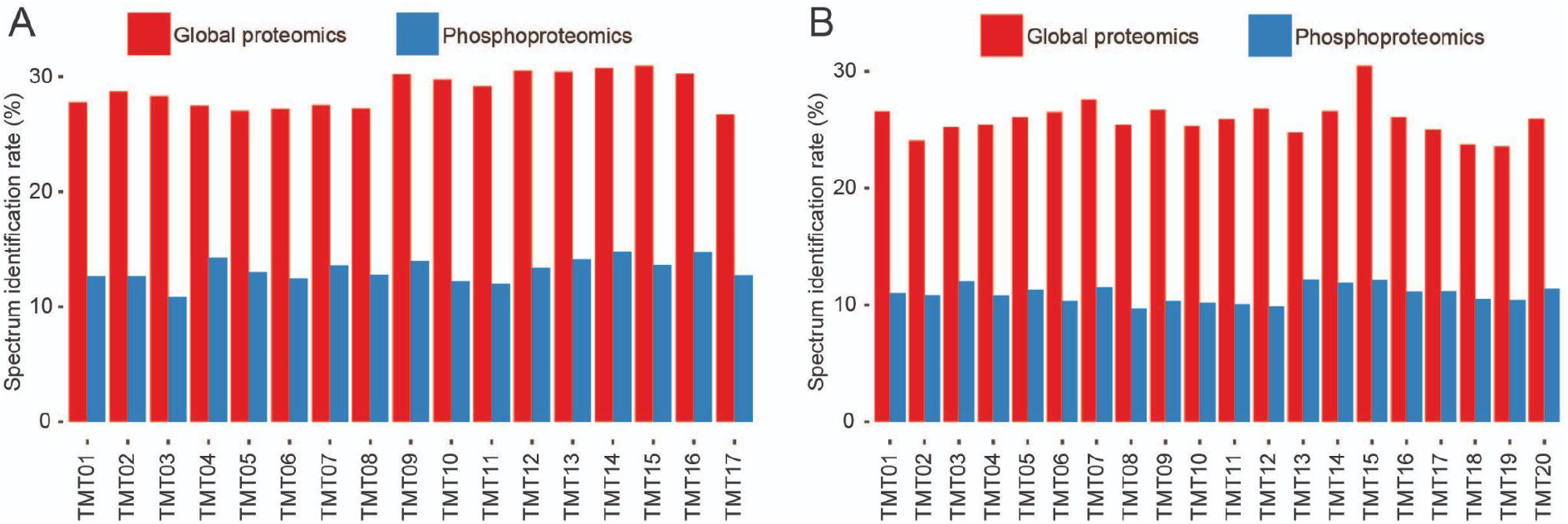
Spectrum identification rate comparison between global proteomics and phosphoproteomics. (A) CPTAC UCEC datasets analyzed by the common data analysis pipeline (CDAP). (B) CPTAC HNSCC datasets analyzed by CDAP.

**Figure S2:**
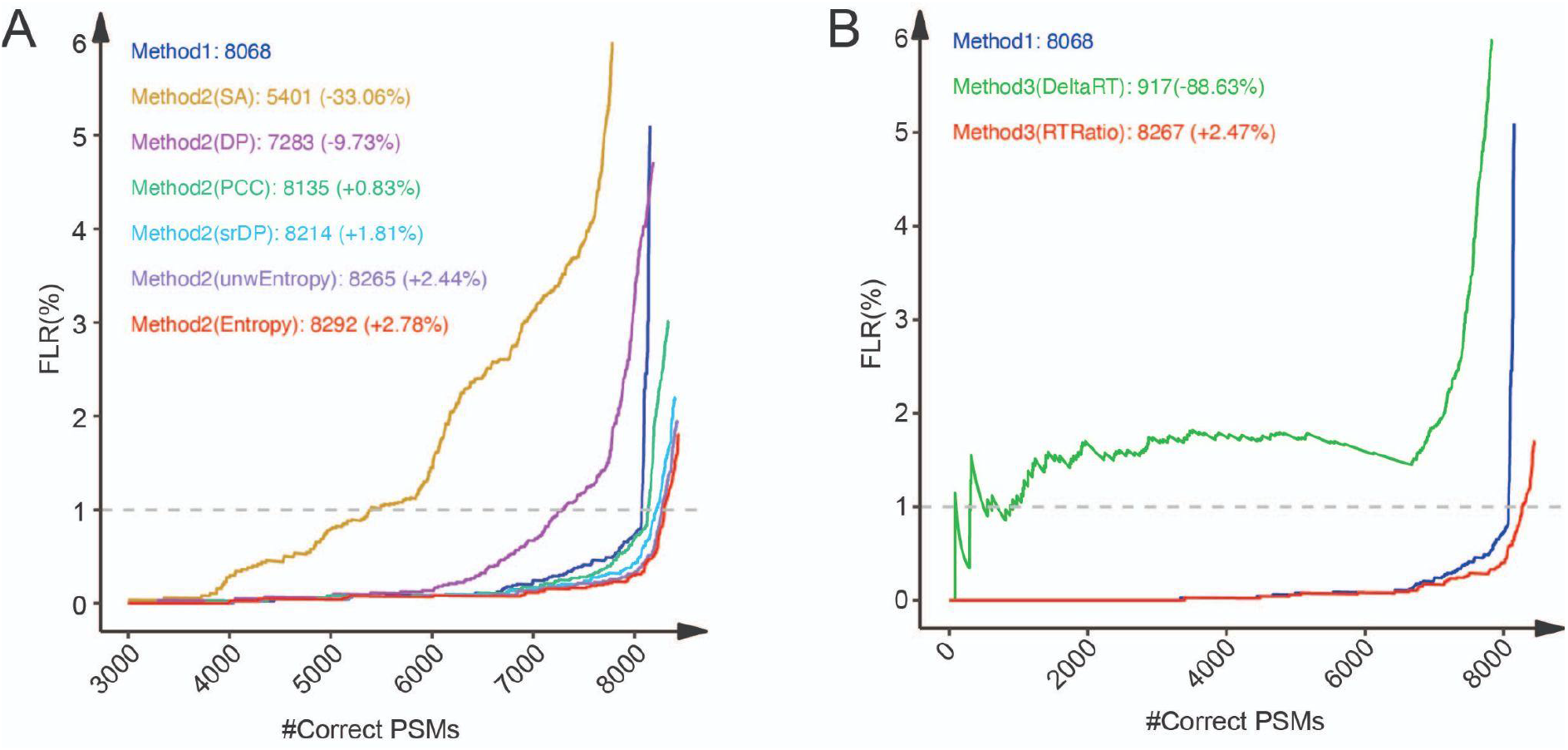
Impact of spectrum similarity (SS) and retention time (RT) difference calculation methods on the performance of site localization on a synthetic phosphopeptide dataset. (A) Impact of six SS calculation methods on the performance of site localization. The number of correctly localized PSMs at different levels of PSM FLR are shown for Method 2 incorporating six different SS calculation methods, respectively. The numbers of correctly localized PSMs at 1% FLR and the percent increase compared with Method 1 (phosphoRS) are indicated. (B) Impact of two RT difference calculation methods on the performance of site localization. The number of correctly localized PSMs at different levels of PSM FLR are shown for Method 3 incorporating two different RT difference calculation methods, respectively. The numbers of correctly localized PSMs at 1% FLR and the percent increase compared with Method 1 (phosphoRS) are indicated.

**Figure S3:**
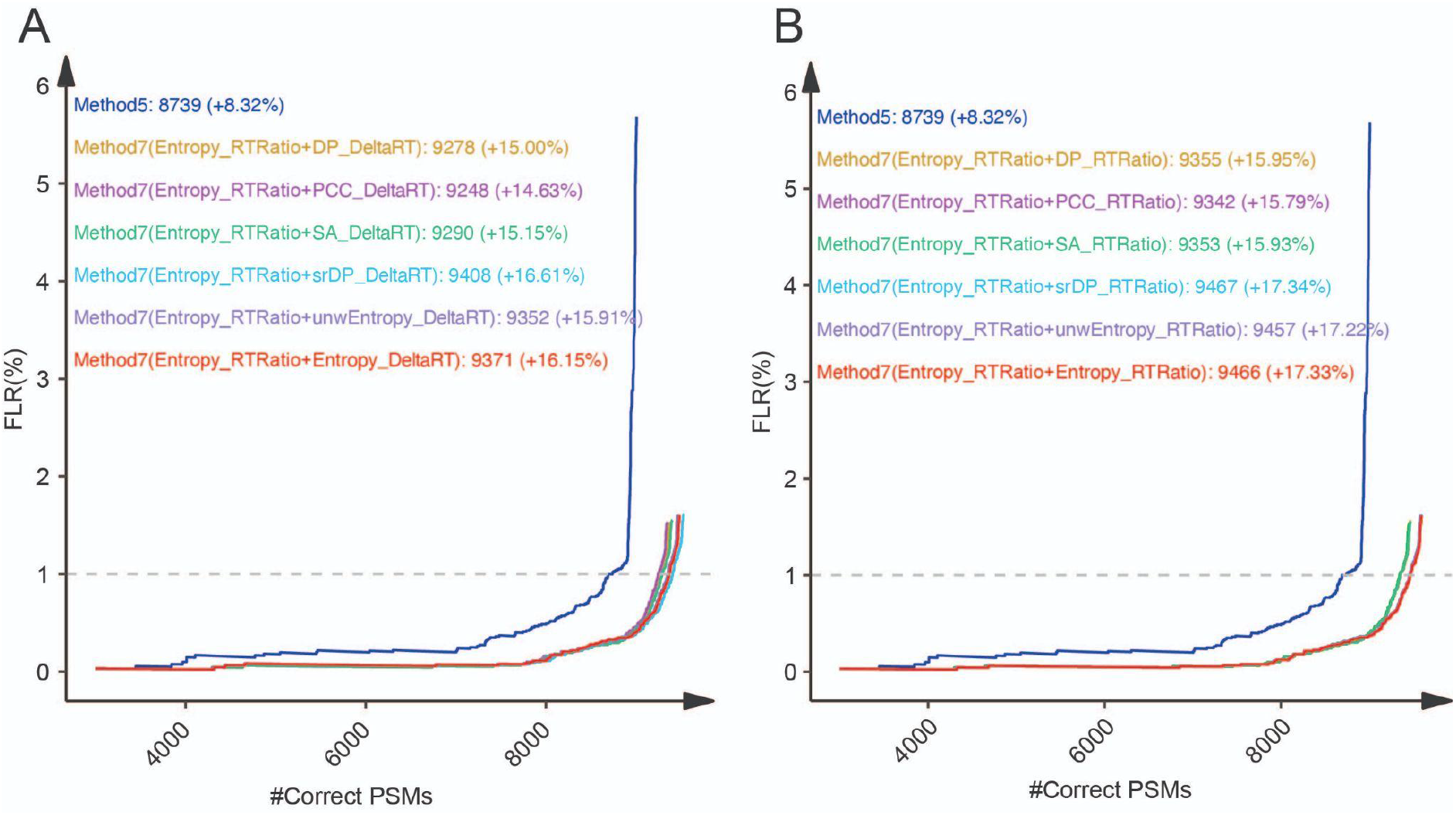
Impact of SS and RT difference calculation methods on the performance of PSM rescoring on a synthetic phosphopeptide dataset. (A) Impact of six SS calculation methods combined with DRT on the performance of PSM rescoring. The number of correctly localized PSMs at different levels of PSM FLR are shown for Method 7 incorporating different SS calculation methods combined with DRT, respectively. The numbers of correctly localized PSMs at 1% FLR and the percent increase compared with Method 5 (phosphoRS+Rescore) are indicated. (B) Impact of six SS calculation methods combined with RTR on the performance of PSM rescoring. The number of correctly localized PSMs at different levels of PSM FLR are shown for Method 7 incorporating different SS calculation methods combined with RTR, respectively. The numbers of correctly localized PSMs at 1% FLR and the percent increase compared with Method 5 (phosphoRS+Rescore) are indicated.

**Figure S4:**
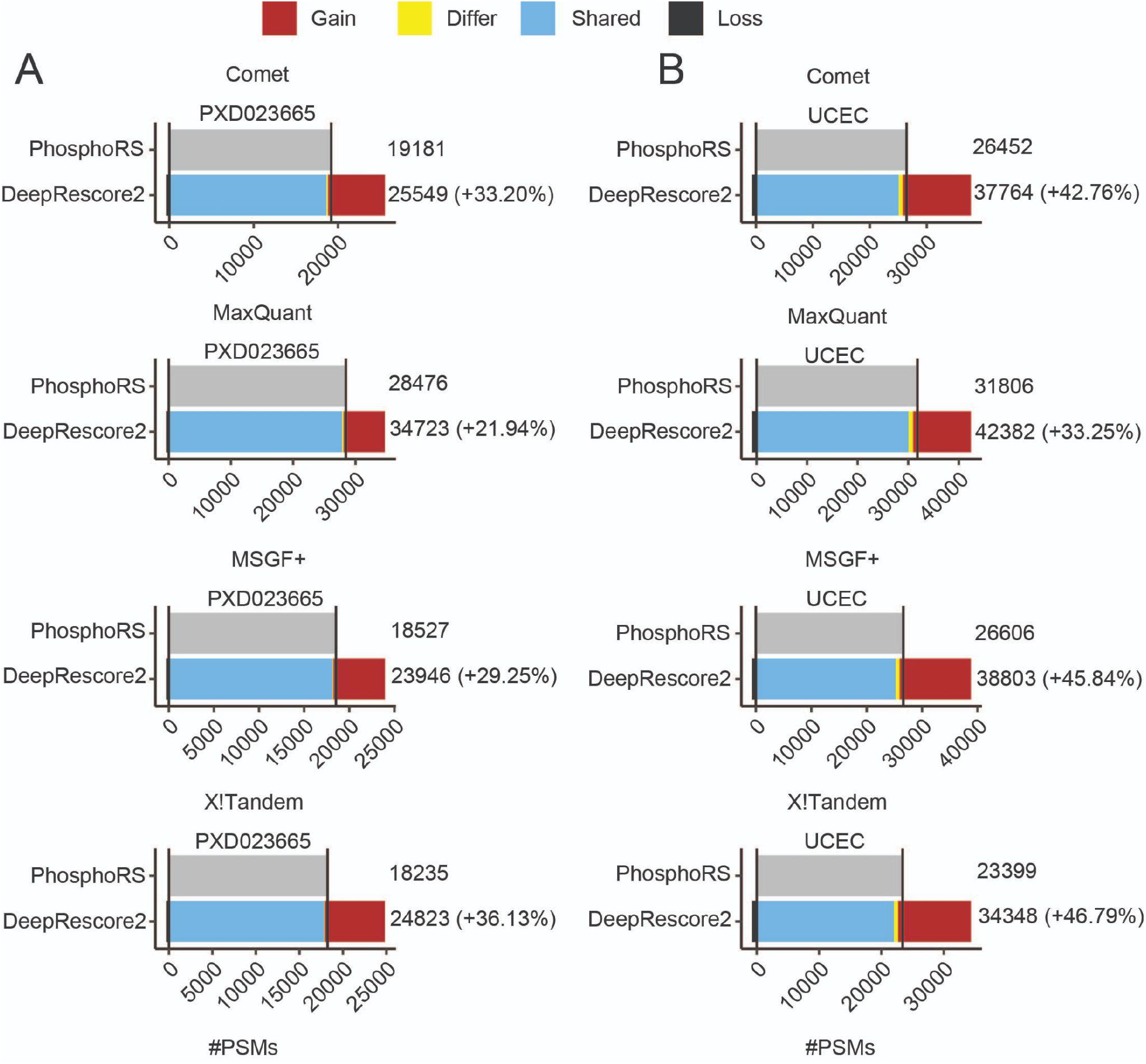
Performance evaluation based on peptide spectrum match (PSM) identifications in two biological datasets using different search engines in combination with PhosphoRS or DeepRescore2. (A) The numbers of PSMs identified from a label-free phosphoproteomic dataset, PXD023665, by four search engines in combination with phosphoRS or DeepRescore2, respectively. (B) The numbers of PSMs identified from the UCEC TMT phosphoproteomic dataset by four search engines in combination with phosphoRS or DeepRescore2, respectively. Gain: PSMs identified by DeepRescore2 but not PhosphoRS. Shared: PSMs identified by both DeepRescore2 and PhosphoRS. Loss: PSMs identified by PhosphoRS but not DeepRescore2.

## Supplementary Tables

**Table S1**. List of datasets included in this study.

**Table S2**. Features used in Percolator in the DeepRescore workflow.

**Table S3**. PSM identifications reported by four different search engines in combination with PhosphoRS or DeepRescore2, respectively.

**Table S4**. Differential expression analysis results and prognosis analysis results based on quantifiable sites reported by PhosphoRS- and DeepRescore2, respectively.

**Table S5**. Prognosis analysis results based on kinase activity inferred from PhosphoRS- and DeepRescore2-derived datasets, respectively.

## Notes

https://github.com/bzhanglab/DeepRescore2

### Competing Interest Statement

The authors have declared no competing interest.

